# Energy efficiency and biological interactions define the core microbiome of deep oligotrophic groundwater

**DOI:** 10.1101/2020.05.24.111179

**Authors:** Maliheh Mehrshad, Margarita Lopez-Fernandez, John Sundh, Emma Bell, Domenico Simone, Moritz Buck, Rizlan Bernier-Latmani, Stefan Bertilsson, Mark Dopson

## Abstract

Extremely oligotrophic deep groundwaters host organisms attuned to the low-end of the bioenergetics spectrum. While all domains of life along with viruses are active in this habitat, the evolutionary and ecological constraints on colonization and niche shifts and their consequences for the microbiome convergence are unknown. Here we provide a comparative genome-resolved analysis of the prokaryotic community in disconnected fracture fluids of the Fennoscandian Shield. The data show that the oligotrophic deep groundwaters flowing in similar lithologies offer fixed niches that are occupied by a common deep groundwater core microbiome. Based on this high resolution “multi-omics” enabled understanding of the underlying mechanisms via functional expression analysis, we conclude that deep groundwater ecosystems foster highly diverse, yet cooperative microbial communities adapted to this setting. The fitness of primary energy producers is increased by ecological traits such as aggregate or biofilm formation. This also facilitates reciprocal promiscuous partnerships with diverse and prevalent epi-bionts, alleviating the “tragedy of common goods”. Hence, instead of a lifestyle where microbes predominantly invest in functions related to maintenance and survival, an episodic and cooperative lifestyle ensures the subsistence of the deep groundwater microbiome. We suggest the name “halt and catch fire” for this way of life.

## Introduction

Prokaryotes are at the base of the food web in deep groundwater habitats that host all domains of life as well as viruses^1,2^. With estimated total abundances of a staggering 5×10^27^cells^3,4^, they are constrained by factors such as bedrock lithology, available electron donors and acceptors, depth, and hydrological isolation from the photosynthesis-fueled surface^4,5^. The limited number of access points to study this environment render our knowledge of deep groundwaters too patchy for robustly addressing eco-evolutionary questions. Consequently, ecological strategies and factors influencing the establishment and propagation of the core deep groundwater microbiome, along with its comprehensive diversity, metabolic context, and adaptations remain elusive.

The deep disconnected biosphere is an environment of constant and frequent selection hurdles, which define not only the composition of the resident community, but more importantly also its strategies to cope with episodic availability of nutrients and reducing agents. In the geochemically-stable and low-energy conditions of the deep biosphere, it is suggested that microbes only occasionally have access to the “basal power requirement” for cell maintenance (e.g., biomass production plus synthesis of biofilms and polymeric saccharides, etc.) or the costly process of duplication^6,7^. Inspecting the expression profile and metabolic context of actively transcribing microbes may reveal the dominant ecological strategies in the deep groundwater and uncover the dimensions of its available niches. The microbial diversity of the terrestrial subsurface has previously been probed by large-scale “omics” in shallow aquifers (Rifle, USA)^8^, a CO_2_ saturated geyser (Crystal Geyser, Utah, USA)^9^, and carbon-rich shales (Marcellus and Utica, USA)^10^. However, extremely oligotrophic and disconnected deep groundwater habitats lack a comprehensive comparative “multi-omics” analysis. The proterozoic crystalline bedrock of the Fennoscandian Shield (~1.8 Ga. years old) hosts two deep tunnels that provide access to disconnected fracture fluids (ca. 170 to 500 meters below sea level, mbsl) running through a similar granite/granodiorite lithology^4,11–14^. The two sites, located in Sweden and Finland, provide a rare opportunity to place the microbiome of deep groundwater under scrutiny (Supplementary FigureS1 depicts the location of large metagenomics datasets for oligotrophic groundwaters).

In this study, we investigate the existence of a common core microbiome and possible community convergence in the extreme and spatially heterogeneous deep groundwater biome. We leverage a large “multi-omics” initiative that combines metagenomes, single cell genomes, and metatranscriptomes from samples collected at two disconnected sites excavated in similar lithology. Through an extensive genome-resolved view and comparative analysis of the communities, we provide strong support for the existence of a common core microbiome in deep groundwaters. The metabolic context and expressed functions of the microbial community were further used to elucidate the ecological and evolutionary processes essential for successfully occupying and propagating in the available niches of this extreme habitat.

## Results and discussion

The Fennoscandian Shield bedrock contains an abundance of fracture zones with different groundwater characteristics; including groundwater source, retention time, chemistry, and connectivity to surface-fed organic compounds. The Äspö HRL and Olkiluoto drillholes were sampled over time, covering a diversity of aquifers representing waters of differing ages plus planktonic versus biofilm-associated communities. In order to provide a genome-resolved view of the Fennoscandian Shield bedrock prokaryotic community, collected samples were used for “multi-omics” integrated analysis by combining metagenomes (*n*=44), single-cell genomes (*n*=564), and metatranscriptomes (*n*=9) (**Supplementary TableS1**). Binning of the 44 generated metagenomes (~1.3 TB sequenced data) resulted in the reconstruction of 1278 metagenome-assembled genomes (MAGs; ≥50% completeness and ≤5% contamination). By augmenting this dataset with 564 sequenced single cell amplified genomes (SAGs; 114 with ≥50% completeness and ≤5% contamination), we present a comprehensive genomic database for the prokaryotic diversity of these oligotrophic deep groundwaters, hereafter referred to as the Fennoscandian Shield genomic database (FSGD; statistics in Figure1A **& Supplementary TableS2**). Phylogenomic reconstruction using reference genomes in the Genome Taxonomy Database (GTDB-TK; release 86) shows that the FSGD MAGs/SAGs span most branches on the prokaryotic tree of life (Figure2). Harboring representatives from 53 prokaryotic phyla (152 Archaeal MAGs/SAGs in 7 phyla and 1240 bacterial MAGs/SAGs in 46 phyla), the FSGD highlights the remarkable diversity of these oligotrophic deep groundwaters. Apart from the exceptional case of a single-species ecosystem developed by *Candidatus* Desulforudis audaxviator in the fracture fluids of an African gold mine^15^, other studies on deep groundwater as well as aquifer sediments have revealed a notable prokaryotic phylogenetic diversity^8,16^. Clustering reconstructed FSGD MAGs/SAGs into operationally defined prokaryotic species (≥95% average nucleotide identity (ANI) and ≥70% coverage) produced 598 genome clusters. Based on the GTDB-TK affiliated taxonomy, a single FSGD cluster may represent a novel phylum, whereas at the lower taxonomic levels, the FSGD harbors genome clusters representing seven novel taxa at class, 58 at order, 123 at family, and 345 at the genus levels. In addition, more than 94% of the reconstructed MAGs/SAGs clusters (*n*=568) represent novel species with no existing representative in public databases (**Supplementary TableS2**). Mapping metagenomic reads against genome clusters represented exclusively by SAGs (*n*=38, Figure 1A) revealed very low abundances for ten genome clusters (16 SAGs), suggesting they might represent rare members of the microbial community in the investigated deep groundwaters (**Supplementary TableS3**).

**Figure 1.**
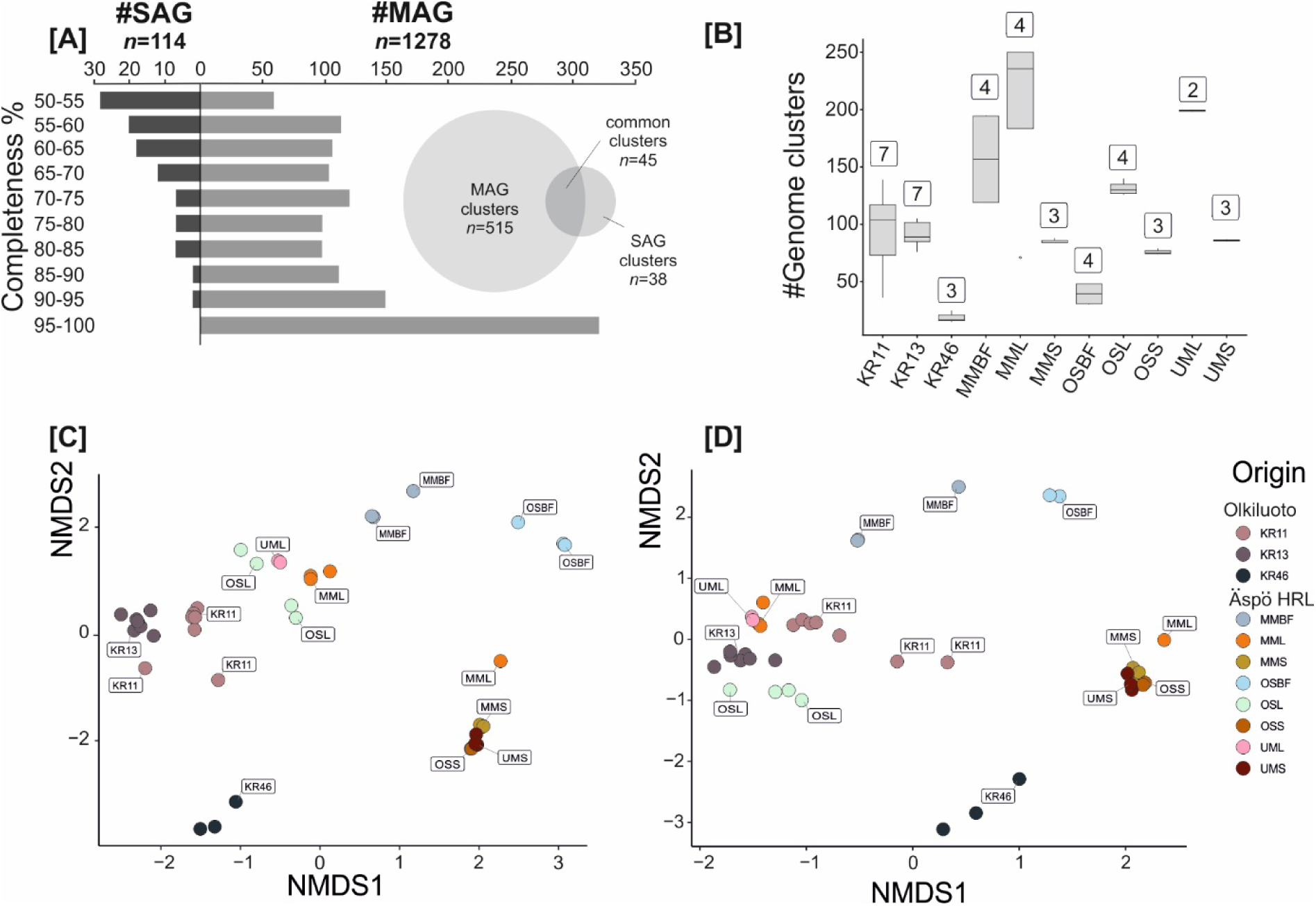
Overview of the FSDB MAGs and SAGs. Statistics of the metagenome assembled genomes (MAGs) and single cell amplified genomes (SAGs) of the Fennoscandian Shield Genomic Database [A]. The species richness as the number of genome clusters present in each borehole/borehole treatment. Numbers on top of each box plot represent the number of metagenomes for each borehole/borehole treatment [B]. NMDS plot of unweighted binary Jaccard beta-diversities of presence/absence of all FSGD reconstructed MAGs/SAGs [C]. MAGs and SAGs belonging to the prevalent clusters present in Äspö HRL and Olkiluoto[D].

**Figure 2.**
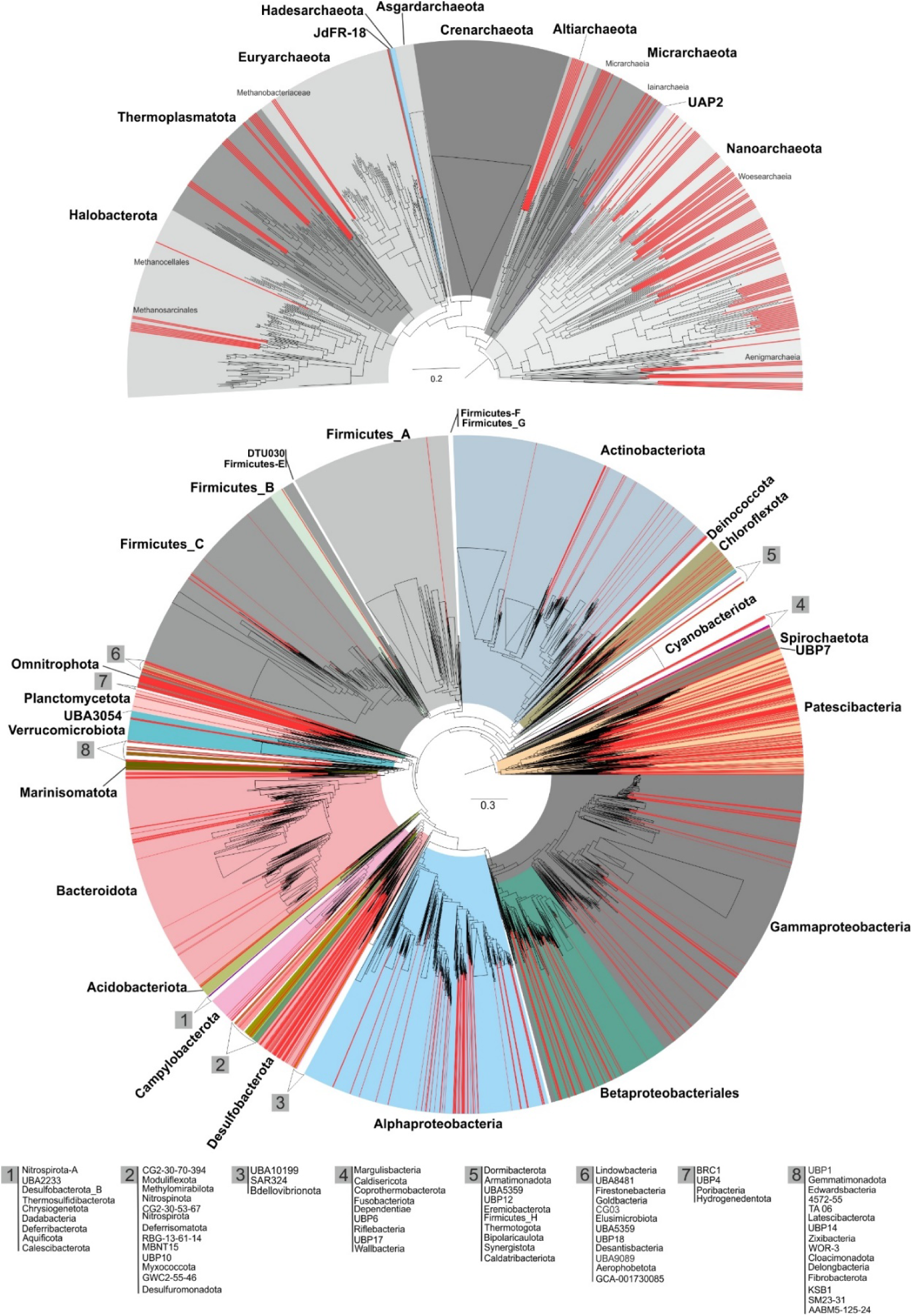
Phylogenetic diversity of reconstructed MAGs and SAGs of the Fennoscandian Shield genomic database (FSGD). Genomes present in genome taxonomy database (GTDB) release 86 were used as reference. Archaea and bacteria phylogeny is represented separately in the top and bottom panels respectively. MAGs and SAGs of the FSGD are highlighted in red.

To explore the community composition of different groundwaters and its temporal variations, presence/absence patterns were computed by competitive mapping of the metagenomics reads against all reconstructed MAGs/SAGs of the FSGD and then normalized for sequencing depth in each metagenome. Since metagenomes were in some cases amplified because of low DNA amounts, we only discuss binary presence/absence values when referring to the community composition to avoid inherent biases in abundance values calculated by counting mapped metagenomics reads. The Äspö HRL metagenomics samples were collected over three years from 2013 to 2016 from five different boreholes. The Olkiluoto metagenomics samples were collected between June and November 2016 from three different drillholes. Communities from each separate borehole cluster together and show only minimal variations in prokaryotic composition over time, hinting at high stability of prokaryotic community composition in the groundwater of the different aquifers. In contrast, the different boreholes feature discrete community compositions (Figures3 & 1C). The observed compositional differences are likely to be at least partially caused by the varying availability of reducing agents and organic carbon in different boreholes, resulting from contrasting retention time, depth, and isolation from surface inputs of organic compounds^14,17^. In the case of Äspö HRL datasets, different treatments (planktonic vs biofilm associated microbes) or size fractions (large vs small fraction) of samples originating from the same boreholes also cluster separately (Figure1C).

Mapping the metagenomic reads against the FSGD identified 340 MAGs/SAGs that were present in groundwater samples from both sites (Figures3 & 1D). These prevalent MAGs/SAGs, cluster in 158 genome clusters taxonomically affiliated to both domain archaea (phyla Nanoarchaeota and Thermoplasmatota; *n*=30) and bacteria (phyla Acidobacteriota, Actinobacteriota, Bacteroidota, Bipolaricaulota, Caldatribacteriota, Campylobacterota, CG03, Chloroflexota, Deferrisomatota, Dependentiae, Desulfobacterota, Firmicutes_A, Myxococcota, Nitrospirota, Omnitrophota, Patescibacteria, Proteobacteria, UBA6262, UBA9089, and Verrucomicrobiota; *n*=310). While the very disconnected nature of the two sites is reflected in their discrete prokaryotic community composition (Figures3 & 1C) the two locations harbor bedrock with similar lithologies. Consequently, they are likely to provide similar niches that may result in convergent species incidence. The presence of the same species in disconnected deep groundwaters, where the bedrock lithology is not the pressing divergence force consolidates the existence of a deep groundwater microbiome primed to occupy the fixed niches available in these habitats. The relatively high phylogenetic diversity of the prevalent species implies a significant role of ecological convergence (due to e.g., availability of nutrient and reducing agents) rather than evolutionary responses. However, this by no means confutes the possibility of an evolutionary convergence as the community clearly undergoes adaptation over the long residence times characteristic for deep groundwaters. For instance, salinity is a proxy for water retention time and it ranges from 0.4 (similar to the brackish Baltic proper) to 1.8% (ca. half that of marine systems) in the different groundwaters (**Supplementary TableS1**). This is reflected in a shift in the isoelectric point of the predicted proteome by decreased prevalence of basic proteins as a potential adaptation strategy in active groundwater microbes to their surrounding matrix over evolutionary timescales (Figure4). The isoelectric point trend towards a reduced prevalence of basic proteins is specifically pronounced in Olkiluoto drillhole OL-KR46 where salinity reaches a maximum of 1.8% (Figure4B). In contrast, the relative frequency of calculated isoelectric points for predicted proteins in metagenomes sequenced from other drillholes in Olkiluoto and Äspö HRL (ranging from 0.4 to 1.2% salinity) are similar to one another, with a relatively higher frequency for basic proteins compared to OL-KR46. The OL-KR46 drillhole provides access to fracture fluids at ~530.6 mbsl where microbial communities have relatively low species richness (15-25 genome clusters per dataset and 27 unique clusters in total), hence representing a distinct community composition from other borehole communities (Figure1B, C, & D). Inspecting the metabolic context of the reconstructed MAGs/SAGs from OL-KR46 suggests a flow of carbon between sulfate-reducing bacteria as the most predominant metabolic guild in the community and acetogens, methanogens, and fermenters as has already been reported^18^. Despite its unique community composition, 85% of the genome clusters represented in the OL-KR46 drillhole are among the prevalent clusters present in groundwaters collected from both sites. These prevalent genomes include representatives of *Pseudodesulfovibrio aespoeensis*^19^, a species originally isolated from 600 mbsl in Äspö groundwater that is present among the FSGD MAGS from Äspö HRL borehole SA1229A and Olkiluoto drillhole OL-KR13 and OL-KR46.

**Figure 3.**
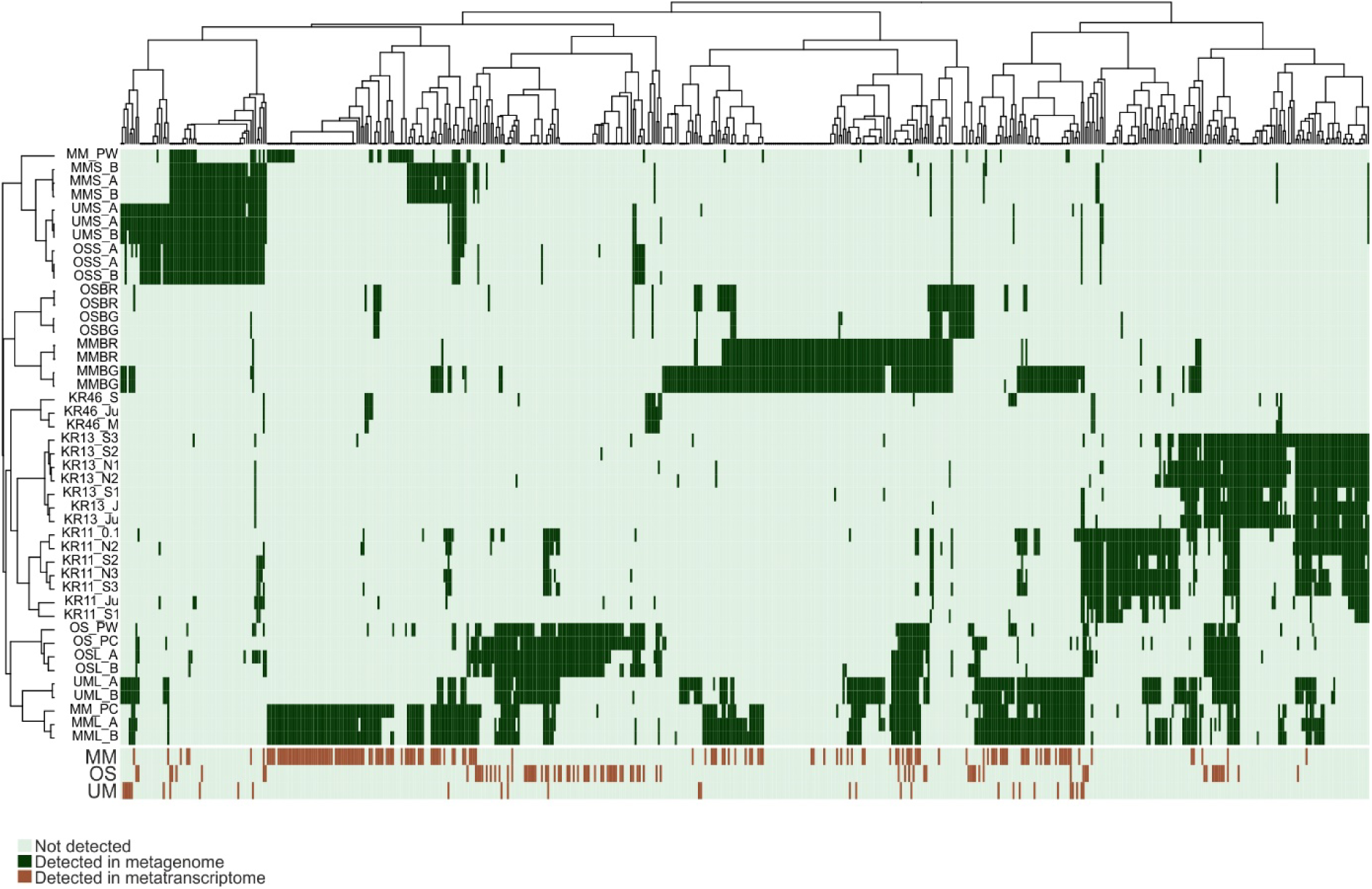
Distribution pattern and transcription status of the FSGD genome clusters along different metagenomes and metatranscriptomes. Each column represents a genome cluster of reconstructed MAGs/SAGs of the deep groundwater datasets of this study. The top heat map depicts the distribution of the representatives of each genome cluster along metagenomics datasets originating from different boreholes of the two Fennoscandian shield deep repositories. The bottom heat map represents the transcription status of the genome cluster representatives in the metatranscriptomes originating from different boreholes of the Äspö HRL. Clusters with all zero values have been removed from the plot (in total 10 clusters that are solely represented by SAGs).

**Figure 4.**
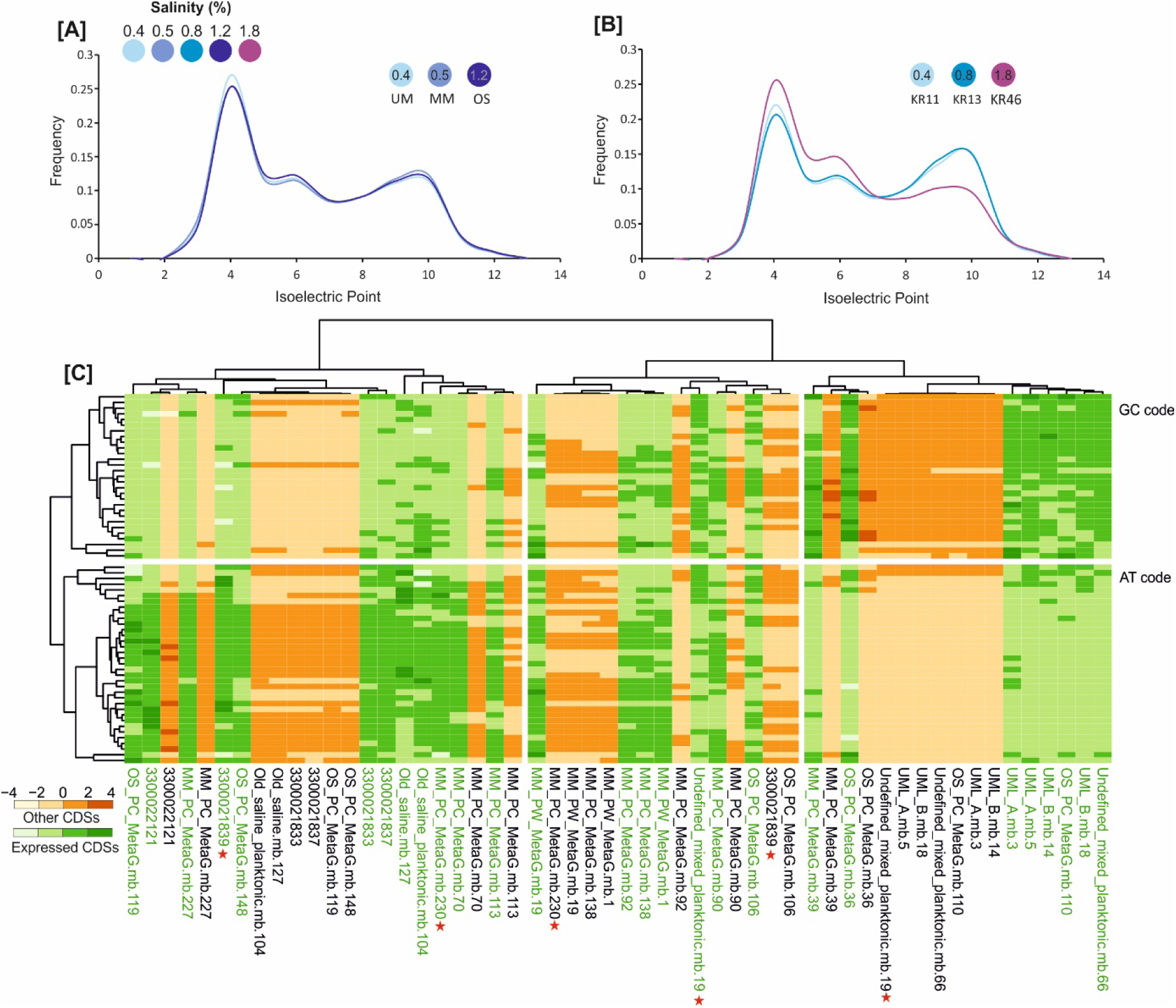
Adaptations of the coding sequences of the deep groundwater microbiota. Relative frequency of isoelectric points in the predicted proteins of assembled metagenomes from Äspö HRL boreholes [A] and Olkiluoto [B]. Salinity of the water flowing in each borehole is shown on the top left legend. Representation of frequency (the expected number of codons, given the input sequences, per 1000 bases) of utilization of synonymous codons across MAGs and SAGs of different genomes clusters of highly expressing Desulfobacterota. Codon usage frequencies are calculated separately for the CDSs for which RNA transcripts are detected (green) and the rest of CDSs not actively being transcribed in the sequenced metatranscriptomes (orange) [C]. Stars are showing cases of potential transcription efficiency control via codon usage bias.

Metabolic context and ecosystem functioning of the resident prokaryotes can provide clues about the features of the fixed deep groundwater niches and whether they are defined predominantly by biotic interactions or abiotic forces. Prior studies prove that representatives of all domains of life are actively transcribing in these deep groundwaters^1,18,20,21^. However, a comprehensive taxonomic and metabolic milieu of the transcribing constituents of the active Fennoscandian Shield community has not been explored. A high-resolution and genome-resolved view of the transcription pattern by mapping metatranscriptomic reads against the FSGD MAGs/SAGs was generated, where actively transcribing genomes, their transcribed genes, and overall metabolic capability was catalogued. This analysis reveals that the majority of FSGD MAGs/SAGs clusters are actively transcribing in the nine sequenced metatranscriptomes derived from the three Äspö HRL boreholes (Figure3). Resources and energy costs dedicated to protein synthesis (i.e., transcription and translation) appear sufficiently large for prokaryotes to be recognized by natural selection and impact their fitness^22,23^. Accordingly, evolutionary adaptations such as codon usage bias and regulatory processes are in place to adjust cellular investments in these processes. Unequal utilization of synonymous codons can have implications for a range of cellular and interactive processes, such as mRNA degradation, translation, and protein folding^24,25^, as well as viral resistance mechanisms, and horizontal gene transfer^26,27^. Calculating the frequency of synonymous codons in the FSGD MAGs/SAGs (Supplementary FigureS2) and those belonging to the highly expressing genome clusters (TPM>10000 arbitrary thresholds; Supplementary FigureS3) reveal variable utilization of synonymous codons in different MAGs/SAGs. These variable patterns are primarily related to the range of GC content (Supplementary FigureS2 & S3). Further exploration of the variable codon utilization among highly expressing representatives of the phylum Desulfobacterota by separate calculation of codon frequency for expressing CDSs (according to the mapping results of metatranscriptomes) highlights cases of potential transcription efficiency controls via codon usage bias in these genomes (Figure4). While in most cases, both expressed CDSs and the rest of CDSs in the genome represent similar synonymous codon frequency and distribution, some genomes display notable differences (e.g. MAGs undefined_mixed_planktonic.mb.19 and MM_PC_MetaG.mb.230 along with SAG 3300021839). The expressed CDSs of these reconstructed genomes encode different functions related to their central role in sulfate reduction metabolism and regulatory functions (**Supplementary TableS5**). Interestingly, the expressed CDSs of the SAG 3300021839 code for heritable host defense functions against phages and foreign DNA (i.e., CRISPR/Cas system-associated proteins Cas8a1 and Cas7). These processes are regulated to be active in case of exposure to virus. Accordingly, the observed differential codon usage frequency of expressed genes versus the rest of genome could hint at its potential role in regulating the efficiency of translation. The same SAG also expresses the ribonuclease toxin, BrnT of type II toxin-antitoxin system. This toxin is known to respond to environmental stressors and cease bacterial growth by rapid attenuation of the protein synthesis most likely via its ribonuclease activity^28^. This expression profile suggests a potential response to phage infection and activation of a host defense system. We suggest that the BrnT/BrnA toxin-antitoxin system could potentially be involved in attenuating the protein synthesis, consequently disrupting the lytic lifecycle of phage.

In deep groundwater habitats, organisms respond to limited energy and nutrient availability by adjusting their energy investment in different expressed traits (see above). Highly expressing cells are presumed to be either equipped with efficient metabolic properties and biotic interactions that are tuned to available niches (including but not limited to the dimensions of fixed niches available to the common core microbiome) or alternatively represent an ephemeral bloom profiting from occasionally available nutrients. To shed light on this, we explored the metabolic context and life style of 86 FSGD genome clusters with high transcription levels (TPM≥10000, arbitrary threshold) comprising 192 MAGs and 35 SAGs. The relatively large phylogenetic diversity of such highly expressing clusters (17 phyla) reaffirms that a considerable fraction of the deep groundwater microbiome has competitive properties with regards to both metabolism and interactions (Figure5).

**Figure 5.**
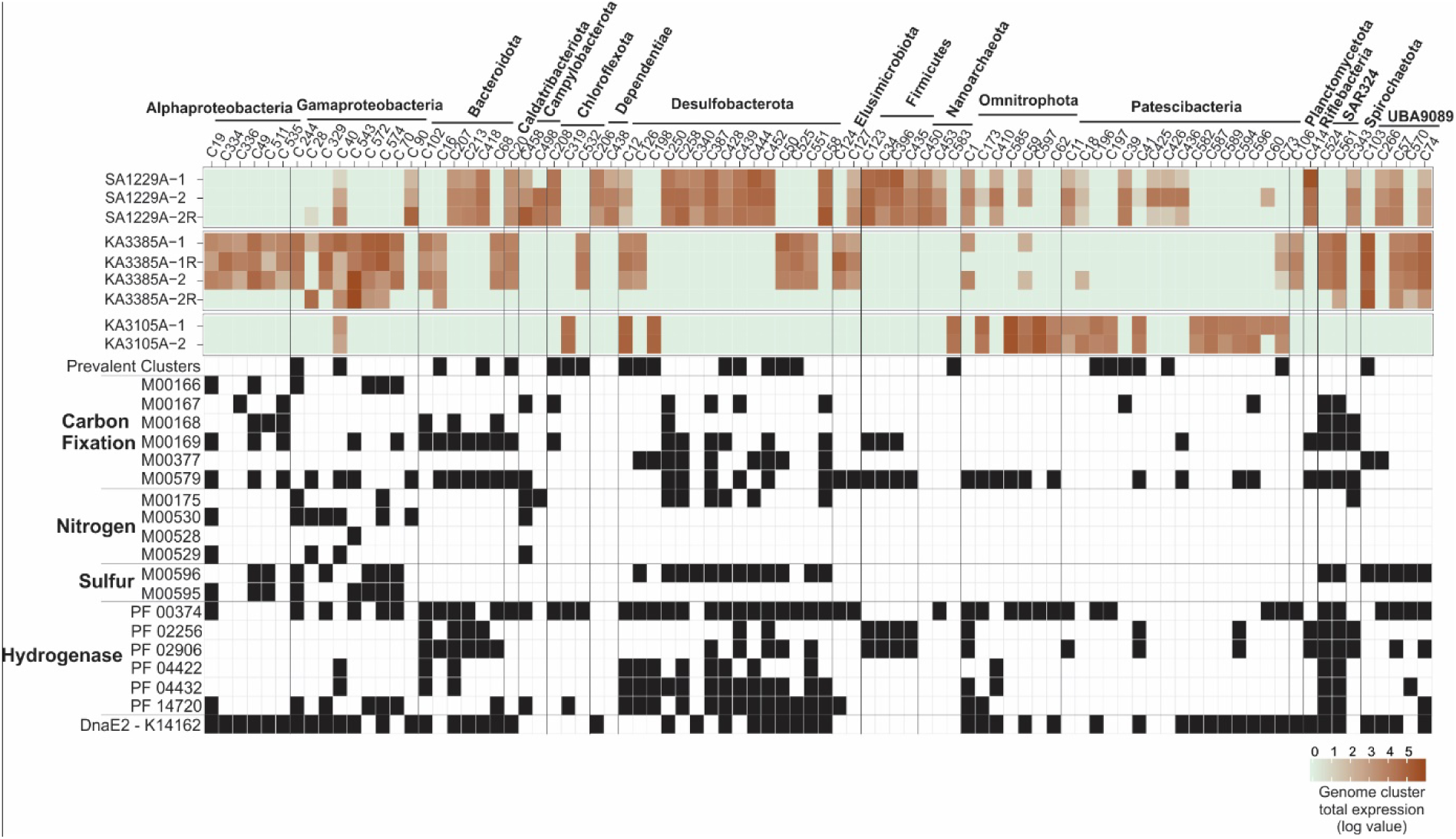
Expression profile and metabolic context of the highly expressing genomic clusters. The expression profile of genome clusters with commutative expression ≥10000 TPM and their metabolic potential for nitrogen and sulfur energy metabolism as well as carbon fixation is represented as KEGG modules presence only if all genes of the module or the key genes of the process are present. Nitrogen fixation (M00175), dissimilatory nitrate reduction (M00530), nitrification (M00528), and denitrification (M00529). Dissimilatory sulfate reduction (M00596) and thiosulfate oxidation by SOX complex (M00595). Reductive pentose phosphate cycle, ribulose-5P => glyceraldehyde-3P (M00166), reductive pentose phosphate cycle, glyceraldehyde-3P => ribulose-5P (M00167), Crassulacean acid metabolism, dark (M00168) and light (M00169), reductive acetyl-CoA pathway (Wood-Ljungdahl pathway) (M00377), and phosphate acetyltransferase-acetate kinase pathway, acetyl-CoA => acetate (M00579). Nickel-dependent hydrogenase (PF00374), iron hydrogenase (PF02256 and PF02906), coenzyme F420 hydrogenase/dehydrogenase (PF04422 and PF04432), and NiFe/NiFeSe hydrogenase (PF14720).

The normalized count of mapped metatranscriptomic reads on genes annotated with functions related to provision of “public-goods” comprise a considerable proportion of the overall transcription profile (ca. 15 to 20% with one case reaching as high as 30%) in deep groundwater metatranscriptomes (Figure6 & **Supplementary TableS6** for a list explored K0 identifiers). One way to alleviate the “tragedy of the commons” imposed by the production and sharing of such “public-goods” is by emergence of local sub-communities either through biofilm formation or aggregation. Such a lifestyle would reduce the number of “cheaters” and provide an evolutionary advantage for cooperation as compared to competition. However, exploitation of “public goods” seems inevitable in groundwaters considering the high phylogenetic diversity and widespread presence of Patescibacteriota representatives (300 MAGs and 19 SAGs forming 152 clusters) and Nanoarchaeota (100 MAGs in 56 clusters). The metabolic context of these reconstructed MAGs/SAGs suggests a primarily heterotrophic and fermentative lifestyle similar to prior reports on the epi-symbiotic representatives of Patescibacteriota and Nanoarchaeota^29^, and accordingly they will be highly dependent on leaky metabolites produced by adjacent cells. The high RNA transcript counts for representatives of these phyla suggests that they have sufficient energy at their disposal to invest in transcription (16 Patescibacteriota and 3 Nanoarchaeota clusters) (Figure5). Their captured transcripts were annotated as ribosomal proteins, cell division proteins (FtsZ), several genes involved in central carbohydrate, lipid, nucleotide, and amino acid metabolism, as well as few genes involved in cofactor and vitamin biosynthesis. To mitigate the burden of “public-goods” providers and in order to stabilize their own syntrophic niche, these epi-symbiotic cells likely participate in reciprocal partnerships where they supply fermentation products (e.g., lactate, acetate, and hydrogen), vitamins, amino acids, and secondary metabolites to their direct or indirect partners in their immediate surroundings. Representatives of these phyla are also detected in the common core microbiome across both groundwater sites (25 Nanoarchaeota MAGs and 81 Patescibacteriota MAGs/SAGs; **Supplementary TableS4**). This implies a significant role of biological interactions in the development of fixed niches in the deep groundwaters. The epi-symbiotic association of Patescibacteriota and Nanoarchaeota with prokaryotic hosts has already been verified for several representatives^30–34^. However, the level and range of host/partner specificity for these associations remain understudied. The incidence of the same genome clusters of Patescibacteriota and Nanoarchaeota representatives in both deep groundwaters, combined with their high expression potential and inferred dependency on epi-symbiotic associations for survival, underscores cooperation as a competent evolutionary strategy in oligotrophic deep groundwaters.

**Figure 6.**
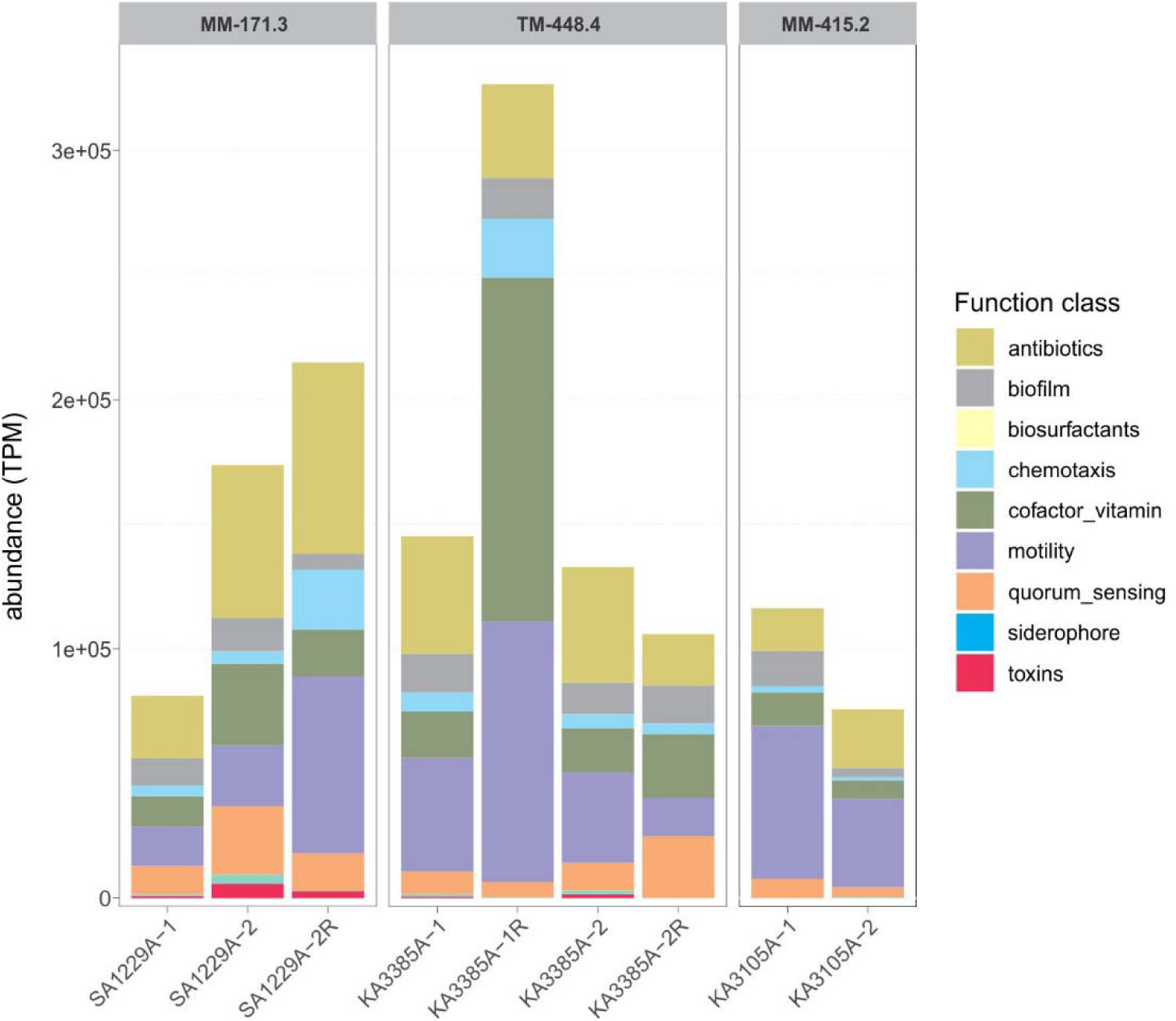
Expression level of functional classes involved in public good provision in the sequenced metatranscriptomes. The list of screened K0 identifiers are shown in **Supplementary Table S6**.

The metabolic overview of FSGD epi-symbiotic MAGs/SAGs suggests these cells to partially depend on the energetic currency provided by their partners/hosts while maintaining partnerships via reciprocal metabolite exchange. Reconstructing the metabolic scheme of highly expressing genome clusters, apart from small heterotrophic cells with a proposed epi-symbiotic lifestyle (Figure5), highlight sulfur as an electron acceptor that is commonly used in the energy metabolism of the deep groundwater microbiome^21^. Dissimilatory sulfur metabolism via the activity of enzymes sulfate adenylyltransferase (*sat*), adenylylsulfate reductase subunits A/B (*aprAB*), and dissimilatory sulfite reductase A/B subunits (*dsrAB*) is present among representative FSGD MAGs/SAGs of domain bacteria (phyla AABM5-125-24, Actinobacteriota, Chloroflexota, Desulfobacterota, Firmicutes-B, Nitrospirota, Planctomycetota, Proteobacteria, SAR324, UBA9089, UBP1, Verrucomicrobiota, and Zixibacteria) and domain archaea (phylum JdFR-18). A total of 203 MAGs and 10 SAGs representing 95 FSGD genome clusters feature the complete gene set for dissimilatory sulfate reduction or at least the *dsrA/B* genes central for the pathway (absence of other genes involved in the pathway could be a consequence of incompleteness). Although the listed taxa have been demonstrated to be capable of sulfite/sulfate reduction^35^, it cannot be ruled out that the *dsrA/B* genes are used in reverse for sulfur oxidation. Representatives of 26 clusters (ca. 30%) of the highly expressing groups represent dissimilatory sulfur metabolism (Figure5). Their complete RNA transcript profile apart from functions related to dissimilatory sulfur metabolism (*aprAB, dsrAB*) are related to a wide range of cellular functions. These include genetic information processing, central carbohydrate turnover, lipid and protein metabolism, biofilm formation, membrane transporters, and other cell maintenance functions as well as genes involved in replication and repair of the genome and cell division. Sulfate-reducing bacteria of the phyla Desulfobacterota and UBA9089 also contain genes encoding for the reductive acetyl-CoA pathway (Wood-Ljungdahl pathway) that is utilized in reverse to compensate for the energy-consuming oxidation of acetate to H_2_ and CO_2_ with energy derived from sulfate reduction^36^. In this process, cells are able to use acetate as a carbon and electron source (Figure5) with various hydrogenase types offering molecular hydrogen as an alternative electron donor^37^.

A notable portion of FSGD transcripts belong to motility related genes (e.g., chemotaxis and flagellar assembly) and studies show that the expression of this costly trait increases in low nutrient environments as an adaptation to anticipate and exploit nutrient gradients^38^. In addition to motility related genes, sulfate-reducing representatives invest in transcription of type IV secretion systems that can facilitate adhesion, biofilm formation, and protein transport. In combination with their expressed chemotaxis genes, we suggest that these motile cells enable cooperation in their local community by attaching to surfaces or by forming aggregates.

Many type II toxin-antitoxin systems (TA) are expressed in the MAGs/SAGs of these oligotrophic groundwaters (e.g., PemK-like, MazF-like, RelE/RelB, and BrnT/BrnA, etc.) likely to alleviate environmental stressors. These molecules are prevalent in prokaryotes^39^ where they participate in a range of cellular processes such as gene regulation, growth arrest, sub-clonal persistence, and cell survival. TA systems often fulfil their regulatory role by halting protein synthesis in response to environmental stimuli. We propose a model in which deep groundwater microbes adjust to the very nutrient-poor conditions while occasionally being flooded by short-lived pulses of nutrients by TA systems triggering bacteriostasis to avoid exhausting the basal energy supply. To restart the cell function in the occasion of ephemeral access to nutrients, the autoregulation of the antitoxin component of the TA system is alleviated to defy the excess of toxin. Finally, we recovered error-prone DNA polymerase (DnaE2) in reconstructed FSGD MAGs/SAGs (Figure5 & Supplementary FigureS4 for the phylogeny of polymerases type-C). These polymerases can be recruited to stalled replication forks and are known to be involved in error-prone DNA damage tolerance^40^ helping with the genome replication We suggest the name “Halt and Catch Fire”^1^ for this mode of subsistence for deep groundwater microbiomes.

## Conclusions

Life in deep groundwaters consisting chiefly of microbes^41^ represents spatial heterogeneity in response to factors like bedrock lithology, depth, and available electron acceptor and donors. Spatial heterogeneity together with limited access points have so far hindered our understanding of the ecological and evolutionary forces governing the colonization and propagation of microbes in the deep groundwater niches. By employing high-resolution exploration of the microbial community in the disconnected fracture fluids running through similar lithologies at two locations, we propose the existence of a common core microbiome in these deep groundwaters. The metabolic context of this common core microbiome proves that both physical filters and biological interactions are involved in defining the dimensions of fixed niches where sulfate-reduction and reciprocal partnerships of epi-bionts are the most competent traits. We suggest an active but “Halt and Catch Fire” strategy for the phylogenetically diverse microbiome of the deep groundwater in response to ephemeral nutrient pulses.

## Methods

### Sampling and multi-omics analysis

Multiple groundwater samples were collected over several years from two deep geological sites excavated in crystalline bedrock of the Fennoscandian Shield. The first is the Swedish Nuclear Fuel and Waste Management Company (SKB) operated Äspö HRL located in the southeast of Sweden (Lat N 57° 26’ 4’’ Lon E 16° 39’ 36’’). The second site is on the island of Olkiluoto, Finland, that will also host a deep geological repository for the final disposal of spent nuclear fuel (Lat N 61° 14’ 31’’, Lon E 21° 29’ 23’’). Water types with various ages and origins were targeted by sampling fracture fluids from different depths. The Äspö HRL samples originated from boreholes SA1229A-1 (171.3 mbsl), KA3105A-4 (415.2 mbsl), KA2198A (294.1 mbsl), KA3385A-1 (448.4 mbsl), and KF0069A01 (454.8 mbsl). The Olkiluoto samples originated from drillholes OL-KR11 (366.7-383.5 mbsl), OL-KR13 (330.5-337.9 mbsl), and OL-KR46 (528.7-531.5 mbsl).

Collected samples were subjected to high-resolution “multi-omics” analysis by combining metagenomics (*n*=27 from the Äspö HRL and n=17 from Olkiluoto), single cell genomics (*n*=564), and metatranscriptomics (*n*=9) (detailed information in **Supplementary Table S1**). Single-cell amplified genomes (SAGs) were captured from KA3105A-4 (*n*=15), KA3385A-1 (*n*=148), SA1229A-1 (*n*=118), OL-KR11 (*n*=138), OL-KR13 (*n*=117), and OL-KR46 (*n*=28) water samples. To probe the expression pattern of the resident community, metatranscriptomic datasets were generated for Äspö HRL samples^1,20^ originating from boreholes KA3105A-4 (*n*=2), KA3385A-1 (*n*=4), and SA1229A-1 (*n*=3). Details of sampling, filtration, DNA/RNA processing, and geochemical parameters of the water samples along with statistics of the metagenomics/metatranscriptomics datasets and SAGs are available in **Supplementary methods** and **Supplementary Table S1**.

### Metagenome assembly

All datasets were separately assembled using MEGAHIT^42^ (v. 1.1.3) (--k-min 21 --k-max 141 --k-step 12 --min-count 2). The datasets originating from the same water type in each location were also processed as co-assemblies in order to increase genome recovery rates (using the same assembly parameters). A complete list of all metagenomic datasets assembled in this study (*n*=44) and the co-assemblies are provided in **Supplementary Table S1**.

### Fennoscandian Shield Genomic Database (FSGD)

The generated “multi-omics” data were used to construct a comprehensive genomic and metatranscriptomic database of the extremely oligotrophic deep groundwaters. Automated binning was performed on assembled ≥2 kb contigs of each assembly using MetaBAT2^43^ (v. 2.12.1) with default settings. Quality and completeness of the reconstructed MAGs and SAGs were estimated with CheckM^44^ (v. 1.0.7). The taxonomy of MAGs/SAGs with ≥50% completeness ≤5% contamination was assigned using GTDB-tk^45^ (v. 0.2.2) that identifies, aligns, and concatenates marker genes in genomes. GTDB-tk then uses these concatenated alignments to place the genomes (using pplacer^46^) into a curated reference tree with subsequent taxonomic classification. Phylogenomic trees of the archaeal and bacterial MAGs and SAGs were also created using the “denovo_wf” subcommand of GTDB-tk (--outgroup_taxon p__Patescibacteria) that utilizes FastTree^47^ (v. 2.1.10) with parameters “-wag -gamma”. Reconstructed MAGs and SAGs were de-replicated using fastANI^48^ (v. 1.1) at ≥95% identity and ≥70% coverage thresholds. A detailed description and genome statistics of the Fennoscandian Shield genomic database (FSGD) is shown in the **Supplementary Table S2** and **Supplementary Methods**.

### Functional analysis of the reconstructed genomes

Annotation of function, computation of isoelectric points and codon usage frequency^49^, abundance, and expression analysis (metatranscriptome) are detailed in the **Supplementary Methods**.

### Data availability

Accession numbers for the metagenomes, SAGs, and metatranscriptomes are provided in **Supplementary Table S1**. The reconstructed genomes of the FSGD are accessible under the NCBI BioProject accession number PRJNA627556. All reconstructed genomes as well as the alignments used for phylogeny reconstruction are deposited to figshare and are publicly available 10.6084/m9.figshare.12170313 and 10.6084/m9.figshare.12170310. all data supporting the findings of this paper are available within this paper and its supplementary material. All the programs used and the version and set thresholds are mentioned in the manuscript and supplementary methods.

## Acknowledgements

The work conducted by the U.S. Department of Energy Joint Genome Institute, a DOE Office of Science User Facility, is supported under Contract No. DE-AC02-05CH11231. The Swedish Research Council (contracts 2018-04311, 2017-04422, and 2014-4398) and The Swedish Nuclear Fuel and Waste Management Company (SKB) supported the study. M.D. thanks the Crafoord Foundation (contracts 20180599 and 20130557), the Nova Center for University Studies, Research and Development, and Familjen Hellmans Stiftelse for financial support. M.D. and D.S. thank the Carl Tryggers Foundation (grant KF16: 18) for financial support. S.B. and M.M. acknowledge financial support from the Swedish Research Council and Science for Life Laboratory. High-throughput sequencing was also carried out at the National Genomics Infrastructure hosted by the Science for Life Laboratory. Bioinformatics analyses were carried out utilizing the Uppsala Multidisciplinary Center for Advanced Computational Science (UPPMAX) at Uppsala University (projects b2013127 and SNIC 2019/3-22) with support from a SciLifeLab-WABI bioinformatics grant.

## Author contributions

M.D., S.B., and R.B.-L. devised the study; M.L.-F. and E.B. collected and processed the samples; M.M., J.S., D.S. and M. B. analyzed the data; M.M. and M.D. interpreted the data and drafted the manuscript; and all authors read and approved the final manuscript.

## Supplementary Figures

**Supplementary Figure S1.**
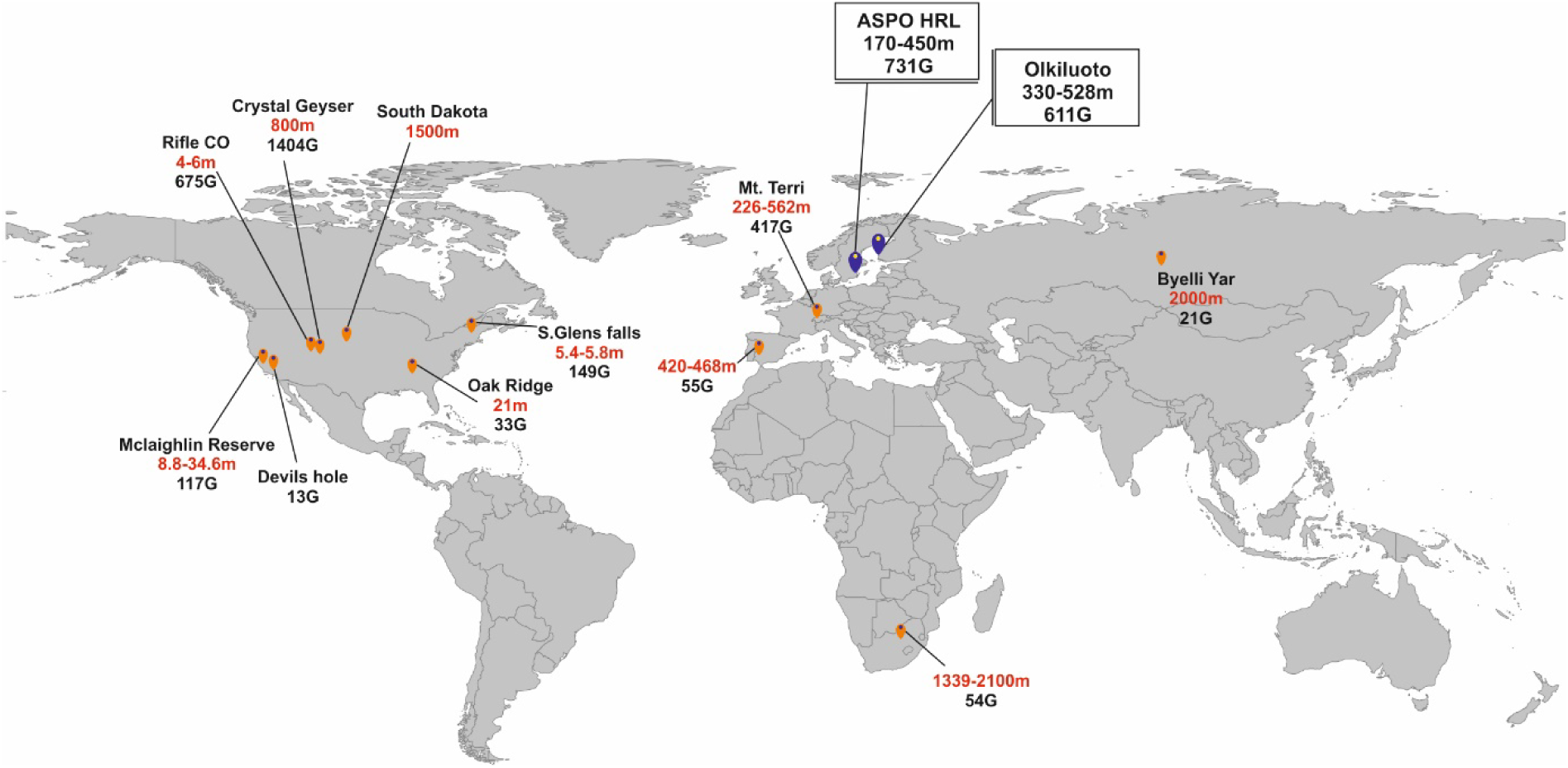
Geographical distribution of publicly available metagenomic datasets sequenced from oligotrophic groundwater samples (landfill groundwater, oil-influenced, and shale samples are not included in this representation). The depth ranges (as mentioned in the corresponding publication) of the samples (in red) and the amount of publicly available sequenced data (presented as Gb) are shown for each location.

**Supplementary Figure S2.**
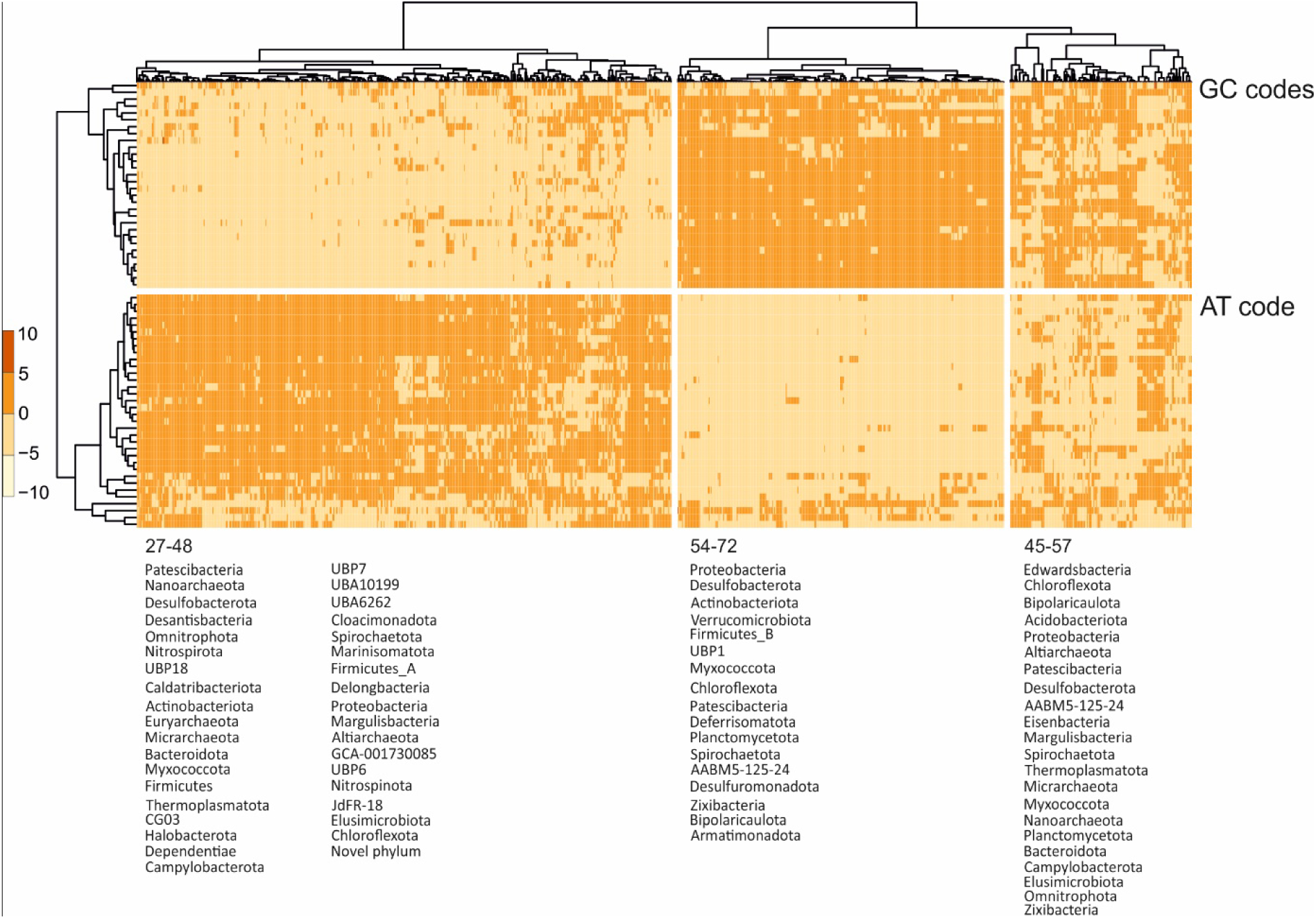
Representation of frequency (the expected number of codons, given the input sequences, per 1000 bases) of utilization of synonymous codons across MAGs and SAGs of the FSGD.

**Supplementary Figure S3.**
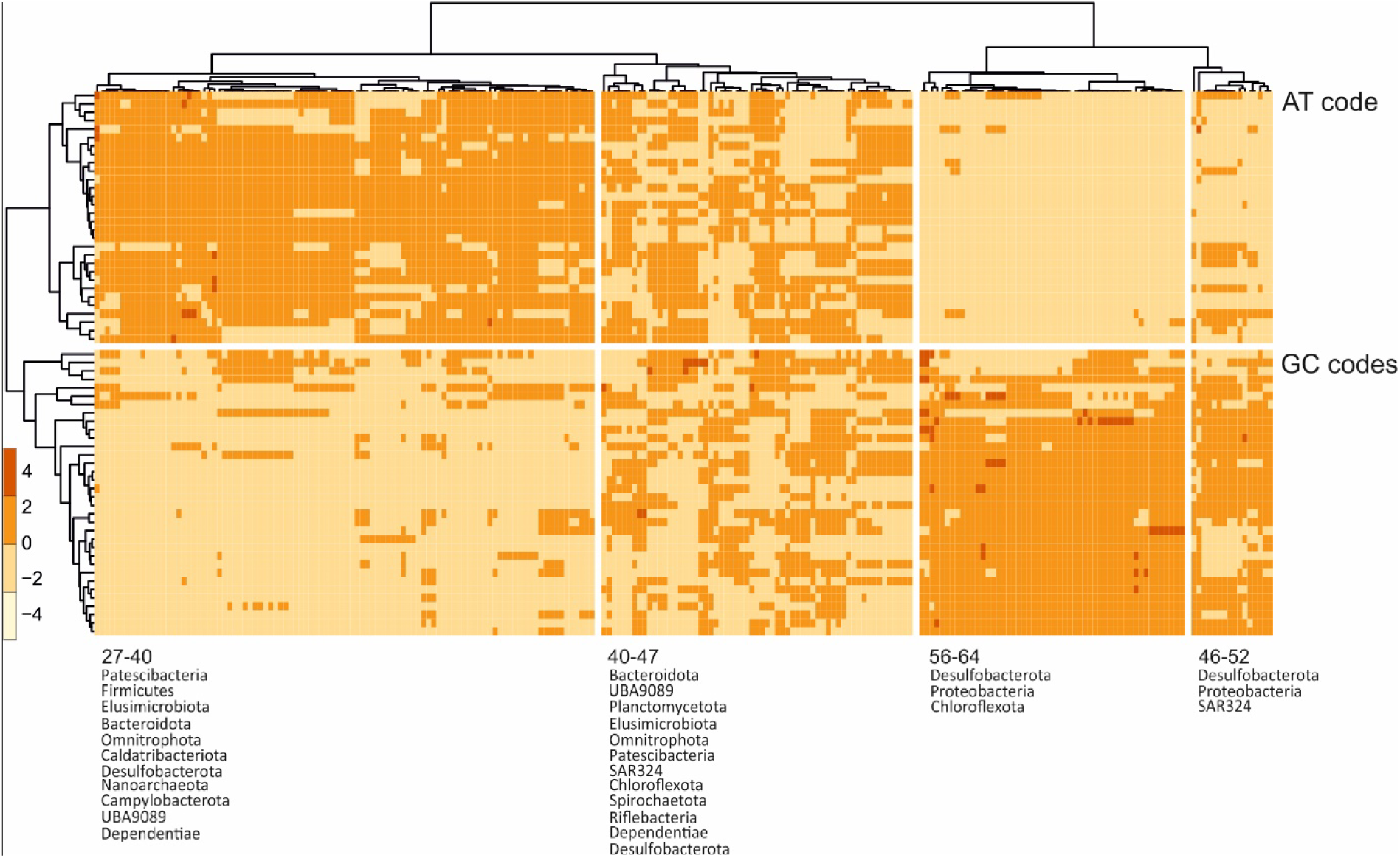
Representation of frequency (the expected number of codons, given the input sequences, per 1000 bases) of utilization of synonymous codons across highly expressed (TPM >10000 arbitrary threshold) MAGs and SAGs of the FSGD.

**Supplementary Figure S4.**
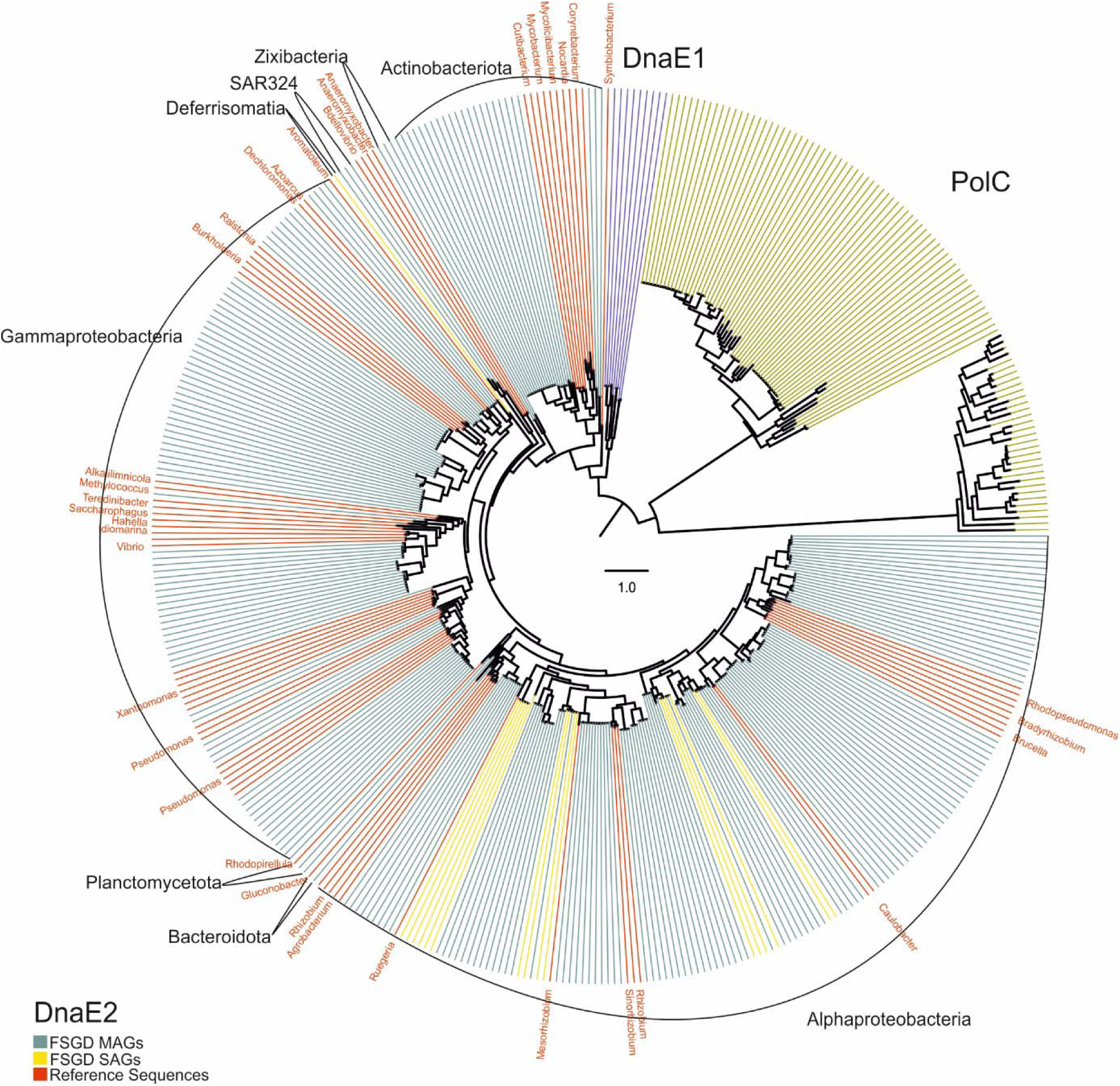
Reconstructed phylogeny of the C-family polymerases. The tree is arbitrarily rooted the PolC branch. Taxonomy of the leafs are written for each group. The reference genomes are the reviewed sequences retrieved from uniprot for each polymerase type (red).

## Footnotes

1 Early operating systems were written in Assembly language that had no built-in safeguards for the hardware. Consequently, mistakes could easily cause components to overheat and be destroyed. “Halt and Catch Fire” was originally an idiom, referring to anything one had done to cause the CPU to fry itself. It got the abbreviation of HCF meaning that the system is fried and the hardware should be replaced. Later, several developers created an HCF three-letter command (or Opcode) that forced the system to halt (without damaging hardware) and it would require a restart to continue working.

