## Supplementary information for "Energy efficiency and biological interactions define the core microbiome of deep oligotrophic groundwater"

**Supplementary Methods**

In this study, the extremely oligotrophic deep groundwaters of two underground research laboratories excavated in the Fennoscandian Shield have been sampled and extensively analyzed via a "multi-omics" approach.

**Site description**

**Äspö Hard Rock Laboratory (Äspö HRL).** This underground laboratory located in the south-east of Sweden (Lat N 57° 26′ 4′′ Lon E 16° 39′ 36′′) is excavated in the Proterozoic crystalline bedrock of the Fennoscandian Shield extending to a depth of 460 m below sea level (mbsl) with 3600 m of total tunnel length[1]. The tunnel was constructed in 1990s and provides access to investigate the microbial life in the deep Fennoscandian Shield groundwater. Along the tunnel, drilled into the bedrock, there are boreholes connecting to water conducting fractures that allow the water to flow towards the tunnel by gravity.

We collected groundwater samples from five different boreholes: SA1229A-1 (171.3 mbsl), KA3105A-4 (415.2 mbsl), KA2198A (294.1 mbsl), KA3385A-1 (448.4 mbsl), and KF0069A01 (454.8 mbsl). These groundwaters carried iron as $Fe^{2+}$, contain dissolved sulfide ($HS^-$), had temporally stable chemistry and $\delta^{18}O$, and neutral pH[2]. However, differential chemical composition and $\delta^{18}O$ values enable us to characterize the origin and age of these groundwaters[3]. Groundwaters SA1229A-1, KA3105A-4, and KA2198A have stable chloride concentrations and $\delta^{18}O$ values, similar to those corresponding to Baltic Sea water[4]. Consequently, these groundwaters have a marine signature and are most likely composed of Baltic Sea water mixed with minor proportions of meteoric water and/or older more saline water residing in the bedrock fractures[5]. The precise infiltration age of this groundwater was unknown, but estimated to be <20 years or even less[6]. They were termed 'modern marine'

(‘MM’) waters, concretely ‘MM-171.3’ for SA1229A-1, ‘MM-294.1’ for KA2198A, and ‘MM-415.2’ for KA3105A-4 groundwaters. Borehole KA3385A-1 contained water with chloride concentrations and $\delta^{18}O$ values in between the groundwaters with saline signature with a very long residence time[7] and marine groundwaters. Therefore, this groundwater was classified as thoroughly mixed (‘TM-448.4’) and is composed of unknown proportions of two or more water types such that the age of this groundwater cannot be assessed[8]. Borehole KF0069A01 had a typical signature of low dissolved organic carbon and other anions, high chloride and sulfate concentrations derived from mineral weathering, an age of millions of years[6], and was defined as ‘old saline’ (‘OS-454.8’). The MM-171.3, MM-415.2, and TM-448.4 boreholes were sampled from November to December 2016 while MM-294.1 and OS-454.8 were both sampled between May to June 2013 (**Supplementary Table S1**).

**Olkiluoto, Finland.** The island of Olkiluoto on the south-west coast of Finland will host a deep geological repository for the final disposal of spent nuclear fuel (Lat N 61° 14’ 31’’, Lon E 21° 29’ 23’’). Groundwater was collected from three drillholes that access fracture fluids at different depths; OL-KR11 (366.7-383.5 mbsl), OL-KR13 (330.5-337.9 mbsl), and OL-KR46 (528.7-531.5 mbsl). Multiple samples were collected during 2016 (OL-KR11 *n* = 7, OL-KR13 *n* = 7 and OL-KR46 *n* = 3). Geochemical parameters of the groundwater were monitored throughout the sampling period and they are available in **Supplementary Table S1**. At Olkiluoto, the groundwater chemistry is stratified with depth. Salinity increases with depth and brackish sulfate-rich groundwater is found up to ~400 m depth, beyond which sulfate-free saline groundwater dominates (Posiva 2013; http://www.posiva.fi/en/databank/posiva_reports/olkiluoto_site_description_2011.1871.xhtml#.XnkY5C2ZNTY). OL-KR11 and OL-KR13 drillholes both access brackish groundwater types (residence time 2,500-8,500 years). Drillhole OL-KR46 accesses a deeper saline groundwater (residence time >10,000 years) (Posiva, 2013).

**Biomass collection, DNA extraction, and metagenome sequencing**

**Äspö HRL.** Planktonic cells were collected after flushing five borehole section volumes on sterile polyvinylidene fluoride (PVDF), hydrophilic, 0.1 µm, 47 mm Durapore membrane filters (Merck Millipore) under *in situ* conditions by connecting a High-Pressure Stainless Steel Filter Holder (Millipore) with a downstream needle valve and pressure gauge directly to the borehole. After filtering an appropriate volume of groundwater, each filter was rolled and placed in a sterile cryogenic tube (Thermo Scientific) and immediately frozen in liquid nitrogen. Samples were frozen at the sampling site to allow transport to the laboratory without any changes in the microbial community. Tubes were stored at -80 °C until further processing. DNA of samples MM-171.3-PC and TM-448.4-PC were extracted using the phenol/chloroform/isoamyl alcohol (24:24:1) method using Phase Lock tubes (Eppendorf). Firstly, 840 µL of TE buffer, pH 8 plus 94 µL of lysozyme (100 mg/mL) were added to each filter before incubation at 37 °C for 30 min. Then, 60 µL of 10 % sodium dodecyl sulfate (SDS) and 6 µL of proteinase K (20 mg/ml) were added, mixed, and incubated at 50 °C for 20 min. Afterwards, an equal volume of phenol/chloroform/isoamyl alcohol was added to the cell lysate, mixed by inverting, and transferred to a Phase Lock Gel tube before centrifugation at 1500 × g for 10 min. Then, another equal volume of phenol/chloroform was added and mixed before centrifuging at 1500 × g for 10 min. The nucleic acid was precipitated by adding an equal volume of ice-cold isopropanol and 0.1 volume of 3M sodium acetate, pH 5.2 and incubating at -20 °C for 60 min. After precipitation, the nucleic acids were centrifuged at 16000 × g and 4 °C for 20 min. The supernatant was discarded and the pellet was rinsed with 500 µL of cold 80 % ethanol. Finally, the pellet was dried at 55 °C on a heat block and re-suspended in 50 µL of TE buffer prior incubation overnight at 4°C. The next day the DNA was incubated at 70 °C for 10 min to help dissolve the last of the DNA. DNA samples termed MM-171.3-PW, MM-415.2-PW, and TM-448.4-PW were extracted using the MO BIO PowerWater DNA isolation kit, following the manufacturer's instructions except that the final DNA re-suspension was performed using 60 µL of eluent[2]. The quality and quantity of the extracted DNA by both methods were analyzed with a Thermo Scientific Nanodrop 2000 and Qubit 2.0 Fluorometer (Life Technologies), respectively. Extracted DNA was stored at -20 °C. Twenty-seven metagenomic datasets were generated from the samples collected from the Äspö HRL. Detailed statistics of the generated metagenomes and the respective sequencing platform are shown in **Supplementary Table S1**. DNA was extracted from the MM-294.1 and OS-454.8 samples as explained in the reference[6].

**Olkiluoto.** To collect biomass for DNA analysis, approximately 10 L of groundwater was pumped directly into a chilled sterile Nalgene filtration unit fitted with a 0.22 µm pore size Isopore polycarbonate membrane (Millipore) and connected to a vacuum pump. After filtration, the membrane filters were rolled and stored in 1.5 mL sterile screwcap tubes. Filters collected for DNA extraction were preserved in 750 mL LifeGuard Soil Preservation Solution (MoBio, Carlsbad, CA, United States) and transferred to the laboratory on dry ice. Filters were stored at -20 °C until further processing. DNA content was extracted using a phenol-chloroform protocol[9] with the following modifications. Firstly, filter pieces were subject to bead-beating (2 × 15 s) prior to incubation in lysozyme for 2 h at 37 °C and secondly, lysate was incubated in Proteinase K (200 mg/mL final concentration) for 2 h. Extracted DNA was measured using the Qubit dsDNA High Sensitivity Assay kit (Thermo Fisher Scientific, Inc.). A total of 17 metagenomic datasets were generated from Olkiluoto, the statistics are shown in **Supplementary Table S1**.

**RNA extraction and metatranscriptome sequencing**

The groundwaters were sampled from the Äspö HRL under *in situ* conditions using two different sampling methods. Firstly, by connecting a sampling device with an in-built fixation system as described in Lopez-Fernandez et al.[10] from June 2015 to March 2016 (**Supplementary Table S1**). Secondly, by connecting a high-pressure stainless steel filter holder (Merck Millipore, USA) with a downstream needle valve and pressure gauge as described in Lopez-Fernandez et al.[8] from September 2015 to January 2016 (**Supplementary Table S1**). In both cases, planktonic cells were collected on sterile hydrophilic polyvinylidene fluoride (PVDF) membranes with 0.1 µm poresize (47 mm Durapore, Merck Millipore, USA) under *in situ* conditions. The cell collection using both sampling methods, RNA extraction, and cDNA generation was performed as previously described[10]. Detailed statistics of the nine sequenced metatranscriptomes are shown in the **Supplementary Table S1**.

**Single cell collection and amplification**

The metagenomics samples were augmented with 564 single-cell amplified genomes (SAGs). The SAGs originate from MM-171.3 (*n*=118), MM-415.2 (*n*=15), and TM-448.4 (*n*=148) borehole samples from the Äspö HRL along with OL-KR11 (*n*=138), OL-KR13 (*n*=117), and OL-KR46 (*n*=28) borehole samples from the Olkiluoto. SAGs were amplified, sequenced, and assembled by the Joint Genome Institute (JGI), USA.

SAGs were de-replicated separately from the MAGs that are containing the exact match of 16S rRNA and those with average nucleotide identity higher than 95% were combined in order to retrieve a higher number of good quality SAGs. These combined SAGs are referred to as several-SAG (s-SAG). A total of 22 SAGs were also sequenced from different water types of

Äspö HRL at the SciLifeLab, Sweden as a pilot study. The SAGs were sequenced using the Illumina platform and assembled using MEGAHIT[11].

SAGs were clustered separately from the MAGs using fastANI (v. 1.1) with 95% average nucleotide identity and 70% minimum overlap. Then, SAGs belonging to the same cluster were analyzed using the 'merge' command in checkm (v. 1.0.7) to find sets of within-cluster SAGs that could potentially be merged in order to increase the completeness. Initially, this resulted in nine pairs of SAGs where the estimated combined genome completeness would increase. GC-profiles were calculated for these 18 SAGs using 1 kbp sliding windows and similar profiles for each pair were validated by manual inspection. Next, redundancies within each pair were investigated by aligning contigs from SAGs with nucmer (v. 3.23) with default settings. Aligned regions were only kept on the longer contigs, by clipping the corresponding stretch from the shorter contigs. If clipping resulted in a contigs <300 bp, that contig was removed completely. In addition, if more than 25% of a contig aligned to another contig, the shorter contig was removed completely. After removing redundant regions, contigs from the SAG pairs were combined. All s-SAGs were again checked for completeness and contamination using Checkm and those with >5% contamination were kept as original SAGs. Detailed information regarding the SAGs is shown in **Supplementary Table S1**.

**Fennoscandian Shield genomic database (FSGD)**

The generated "multi-omics" data were used to construct a comprehensive genomic and metatranscriptomic database of different water-types of the extremely oligotrophic deep groundwater.

The sequenced metagenomes were quality checked and trimmed using Trimmomatic[12] (v. 0.36) with settings to trim the Illumina TruSeq adapter ('TruSeq3-PE-2.fa:2:30:15

LEADING:3 TRAILING:3 SLIDINGWINDOW:4:15 MINLEN:31'). For the six samples sequenced on the MiSeq platform, reads were first cropped to 125 bp by trimming the right end of the reads prior to adapter and quality trimming as above. This was done to recover more paired-end reads from these samples that all had lower quality bases at the end of reverse reads. Each dataset was assembled separately as well as co-assemblies on those datasets originating from the same water type in each sampling site using MEGAHIT[11] (v. 2.12.1) with settings (--k-min 21 --k-max 141 --k-step 12 --min-count 2). Contigs ≥2kb in each assembly were automatically binned using metabat2[13] with default setting. CheckM[14] was used to estimate the genome completeness of MAGs, SAGs, and s-SAGs. Those with completeness ≥50% and contamination ≤5% were considered for down-stream analysis. In addition, SAGs with <50% completeness were considered for down-stream analysis if they clustered with another SAG in the SAG-specific fastANI step (see above) and if they had a genome size ≥500 kbp.

**Genome de-replication.** MAGs and SAGs were clustered using fastANI[15] (v. 1.1) at ≥95% identity and ≥70% coverage threshold. Those genomes in a single cluster are considered as representatives of a single population.

**Genome taxonomy and phylogeny.** Taxonomic affiliation of the FSGD MAGs/SAGs was assigned using GTDB-tk (v. 0.2.2) with reference to the release 86 database[16]. The pplacer alignments generated by the GTDB-tk for bacteria and archaea were curated and used for phylogeny reconstruction using FastTree[17] (v. 2.1.10) with parameters '-wag -gamma'.

**Gene annotation and functional analysis.** Prodigal[18] (v. 2.6.2) was run in metagenomic mode ('-p meta') for predicting protein-coding genes in the assembled contigs. This was followed by functional annotation of the predicted proteins using eggnog-mapper[19] (v. 2.2.1) with the eggnog_5.0 database, and pfam_scan.pl (v. 1.6) with the 31.0 release of the PFAM database. Reconstructed MAGs and SAGs were initially annotated using Prokka[20] (v. 1.12) followed by further annotation with eggnog-mapper and pfam_scan.pl using the same databases as for the metagenomic assemblies. Enzyme EC numbers, and KEGG orthologs, pathways and modules were assigned from the eggnog-mapper output.

**Genome abundance.** Metagenomics reads were mapped against all MAGs/SAGs that passed the criteria for downstream analysis using bowtie2[21] (v. 2.3.3.1) with parameters '--very-sensitive --no-unal'. This was followed by removal of duplicates using MarkDuplicates from the picard suite (v. 2.18.6). Mapped reads were counted using featureCounts[22] (v. 1.6.1) with settings '-M -B' to count multi-mapping and only count read-pairs with both ends aligned. The raw counts were normalized as transcripts per million (TPM) in order to calculate MAGs/SAGs abundance in each metagenome. Based on the calculated TPM the MAGs/SAGs were considered detected in the metagenome if they show value ≥1 and not detected if the value is <1.

**Computation of Isoelectric point and codon usage frequency.** The isoelectric point calculation for the protein sequences as well as amino acid features were calculated using pepstat software in the EMBOSS package[23]. The codon usage frequency of the coding regions was calculates using software cusp in the EMBOSS package[23].

**Gene phylogeny.** The phylogeny of the C-family polymerases was reconstructed by using the reference genomes are the reviewed sequences retrieved from uniprot for each polymerase type. The annotation of the protein coding sequences annotated as DnaE2 was verified by using this phylogeny. Sequences were aligned using Kalign[24] (2.04) and FastTree was used for creating the maximum likelihood tree (JTT +CAT model, gamma approximation).

**Metatranscriptome analysis.** The sequenced metatranscriptomes were quality checked and trimmed using Trimmomatic[12]. The rRNA reads were filtered out using cmsearch[25]. The remaining reads were mapped against the FSGD MAGs/SAGs. The expressed genetic content of each MAG/SAG were extracted at the threshold of 100 TPM and the pattern of gene expression and the expressed content of each MAG/SAG was analyzed.
